# Somnotate: A probabilistic sleep stage classifier for studying vigilance state transitions

**DOI:** 10.1101/2021.10.06.463356

**Authors:** Paul J. N. Brodersen, Hannah Alfonsa, Lukas B. Krone, Cristina Blanco Duque, Angus S. Fisk, Sarah J. Flaherty, Mathilde C. C. Guillaumin, Yi-Ge Huang, Martin C. Kahn, Laura E. McKillop, Linus Milinski, Lewis Taylor, Christopher W. Thomas, Tomoko Yamagata, Russell G. Foster, Vladyslav V. Vyazovskiy, Colin J. Akerman

## Abstract

Electrophysiological recordings from freely behaving animals are a widespread and powerful mode of investigation in sleep research. These recordings generate large amounts of data that require sleep stage annotation (polysomnography), in which the data is parcellated according to three vigilance states: awake, rapid eye movement (REM) sleep, and non-REM (NREM) sleep. Manual and computational annotation methods currently ignore intermediate states because the classification features become ambiguous. However, these intermediate states contain important information regarding vigilance state dynamics. Here, we present a new classifier, “Somnotate”, which produces automated annotation accuracies that exceed human expert performance on mouse electrophysiological data, is robust to errors in the training data, compatible with different recording configurations, and maintains high performance during experimental interventions. Somnotate is a probabilistic classifier based on a combination of linear discriminant analysis (LDA) with a hidden Markov model (HMM). A unique feature of Somnotate is that it quantifies and reports the certainty of its annotations, enabling the experimenter to identify ambiguous recording periods in a principled manner. We leverage this feature to identify epochs that exhibit intermediate vigilance states, revealing that many of these cluster around state transitions, whereas others correspond to failed attempts to transition. We show that the success rates of different transitions can be experimentally manipulated and explain previously observed sleep patterns. Somnotate can thus facilitate the study of sleep stage transitions and offers new insight into the mechanisms underlying sleep-wake dynamics.

**Author summary:** Typically, the three different vigilance states – awake, REM sleep, and non-REM sleep – exhibit distinct features that are readily recognised in electrophysiological recordings. However, particularly around vigilance state transitions, epochs often exhibit features from more than one state. These intermediate vigilance states pose challenges for existing manual and automated classification methods, and are hence often ignored. Here, we present ‘Somnotate’ - an open-source, highly accurate and robust sleep stage classifier, which supports research into intermediate states and sleep stage dynamics. Somnotate quantifies and reports the certainty of its annotations, enabling the experimenter to identify abnormal epochs in a principled manner. We use this feature to identify intermediate states and to detect unsuccessful attempts to switch between vigilance states. This provides new insights into the mechanisms of vigilance state transitions in mice, and creates new opportunities for future experiments.

## Introduction

Long-term electrophysiological recordings from freely behaving mice and other laboratory animals are a popular and powerful mode of investigation for neuroscientists, particularly sleep researchers (1–3). This approach affords the study of a wide range of animal behaviours and associated neurophysiological activities, under experimentally controlled conditions. The recordings typically incorporate an electroencephalogram (EEG) signal recorded from the cortical surface at one or more locations, and may also include electromyogram (EMG) recordings from relevant muscle groups. As the vigilance state profoundly affects the behaviour and physiology of the animal, the first step in the analysis is typically sleep stage annotation. This involves the parcellation of the data into three vigilance states: awake, rapid eye movement (REM) sleep, and non-REM (NREM) sleep.

Whilst sleep stage annotation is typically performed by human experts, there are two principal motives for developing effective automated methods for annotating sleep data. First, manual annotation is time-consuming, as an experienced scorer typically requires several hours to annotate a single 24-hour data set. This places a significant burden on the analysis stage of most experiments, meaning that automation is required to conduct experiments at a scale that would be otherwise difficult to imagine (4,5). To this end, several automated methods have been developed for sleep stage annotation, primarily in the context of human clinical data (6–17), but also in the context of laboratory animals (18–26).

The second principal motive for developing automated sleep stage annotation is that the underlying electrophysiological signals can be ambiguous with respect to the sleep stage. This is especially the case during intermediate sleep states, when EEG signals can exhibit features of more than one vigilance state. Local slow wave activity for example, which is normally considered a hallmark of NREM sleep, has been observed during REM and awake states across different cortical regions in humans (27,28) and rodents (29–31). When faced with such ambiguity, manual annotations often differ between scorers. Automated methods remove such inter-rater variance and afford new opportunities to systematically describe these intermediate states, and investigate their importance for sleep-wake dynamics.

Here, we present ‘Somnotate’ - an open-source, highly accurate and robust sleep stage classifier, which supports research into intermediate states and sleep stage dynamics. Somnotate is shown to consistently exceed the accuracy of expert manual annotations on rodent electrophysiological data and is remarkably robust to errors in the training data and across a series of experimental manipulations. Somnotate is a probabistic classifier that quantifies the certainty of its predictions, thereby identifying epochs that present abnormal electrophysiological signals. This not only affords the opportunity to review the algorithm’s predictions, but also offers a principled method for describing intermediate sleep states across large data sets, where an exhaustive review of the data would be impractical. Here, we show that intermediate states are typically present around vigilance state transitions, but also identify failed transition attempts. We show that different types of state transitions have markedly different success rates, which may explain commonly observed sleep patterns, and we demonstrate that the success rates are altered by interventions such as sleep deprivation. Somnotate can thus facilitate the study of sleep stage transitions and offers new insight into the mechanisms underlying sleep-wake dynamics.

## Results

### Establishing a probabilistic vigilance state classifier

In machine learning, classifiers typically learn a transformation that maps a sample consisting of a set of input values or features, onto a single output value or category. This forms the basis of most automated methods for sleep stage classification, such as decision trees, linear discriminant analysis (LDA), support vector machines, and neural networks (10,20,22–24,32). A key weakness of deterministic machine learning approaches is that each input is mapped onto a single output. As a result, any ambiguity in the sample is ignored, and there is no indication how certain the classifier is in its estimate of the vigilance state. This problem can be solved by using a probabilistic classifier that maps each sample onto a probability distribution over vigilance states. The simplest probabilistic classifier is the naive Bayes classifier, which predicts the most likely state ŝ ∈ S of each sample d ∈D in the test set based on the probability of the sample given each state P(D|S) weighted by the frequency of states P(S):

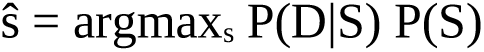

A shortcoming of this approach is that each sample is classified independently and, as the input data can be noisy, this can result in misclassifications and an overestimation of the number of state transitions. Human scorers avoid these issues by using contextual information, subjectively integrating the evidence within each time bin, with an estimate of the likelihood of each state based on the broader context; the high performance of recurrent neural networks compared to other methods is likely due to similar reasons. Algorithmically, the simplest way to integrate predictions based on individual samples with contextual information is to smooth the inferred state sequence by determining the most common vigilance state surrounding the sample of interest. However, vigilance states can be short-lived, which poses a difficult problem for such an approach. For example, mice often transition from REM sleep to NREM sleep via a brief period in which their EEG and EMG activity reflect the awake state (33–36). A probabilistic classifier that can systematically integrate contextual information without smoothing is the hidden Markov model (HMM). The HMM classifier is conceptually similar to the naive Bayes classifier, with the difference being that the prior probability of the state, P(S), is not approximated by its expected frequency, but rather depends on the overall most likely state sequence given the context of the surrounding samples.

A drawback of probabilistic classifiers is that they require an estimate of the multivariate probability distribution over input values for each state. With each additional feature, the number of samples needed to accurately estimate these probability distributions increases exponentially. Consequently, HMMs that have been used in vigilance state annotation previously used low dimensional, hand-crafted features (37–39) or state sequences inferred by other algorithms (7,8,13). Due to the biases of human perception, manual feature engineering from complex signals carries the risk of discarding information that might be valuable for a downstream machine learning classifier. To avoid this, we set out to combine probabilistic classifiers with LDA, which can be trained to automatically extract low dimensional features from complex, high-dimensional inputs, whilst retaining the maximal amount of (linearly decodable) information about the target classes.

To benchmark the performance of our new approach, and to illustrate the advantage conferred by incorporating contextual information, we compared the performance of an HMM to a naive Bayes classifier and an LDA classifier on a data set of six 24-hour (i.e. 144 hours total) simultaneous recordings of anterior EEG, posterior EEG, and EMG in freely behaving mice (Materials and methods), which were independently scored by at least four experienced sleep researchers to annotate awake, NREM, and REM states (**Fig 1A**). For all classifiers, the data was prepared in the same way. The EEG and EMG traces (**Fig 1B**) were subsampled to 256 Hz and converted to multitaper spectrograms (40). The power values in each frequency band were normalised by applying a log(x + 1) transformation and then converting the result to z-scores (**Fig 1C**). As power is exponentially distributed in EEG and EMG signals, a log(x + 1) transformation results in approximately normally distributed values. This facilitates the determination of robust linear discriminants and the targeted dimensionality reduction via linear discriminant analysis (LDA; **Fig 1D**), the final step. Thresholding the LDA representations yielded the LDA classification. For the naive Bayes classifier, multivariate Gaussian distributions (one for each state; **Fig 1G**) were fitted to the low dimensional representation of the samples in the training data set, and the corrsponding state annotations were used to determine the expected frequency of each state (**Fig 1H**). The states corresponding to samples in the test set were then predicted based on the probability of the sample given each state P(D|S) (**Fig 1E**), weighted by their frequency P(S). For the HMM, the probability given each state P(D|S) was computed in the same way as for the Bayes classifier. However, based on the expected transition frequencies observed in the state sequences in the training data set (**Fig 1I**), P(S) was computed using the Baum-Welch algorithm (**Fig 1F)** and the most likely state sequence through the test set was computed using the Viterbi algorithm (41).

**Fig 1.**
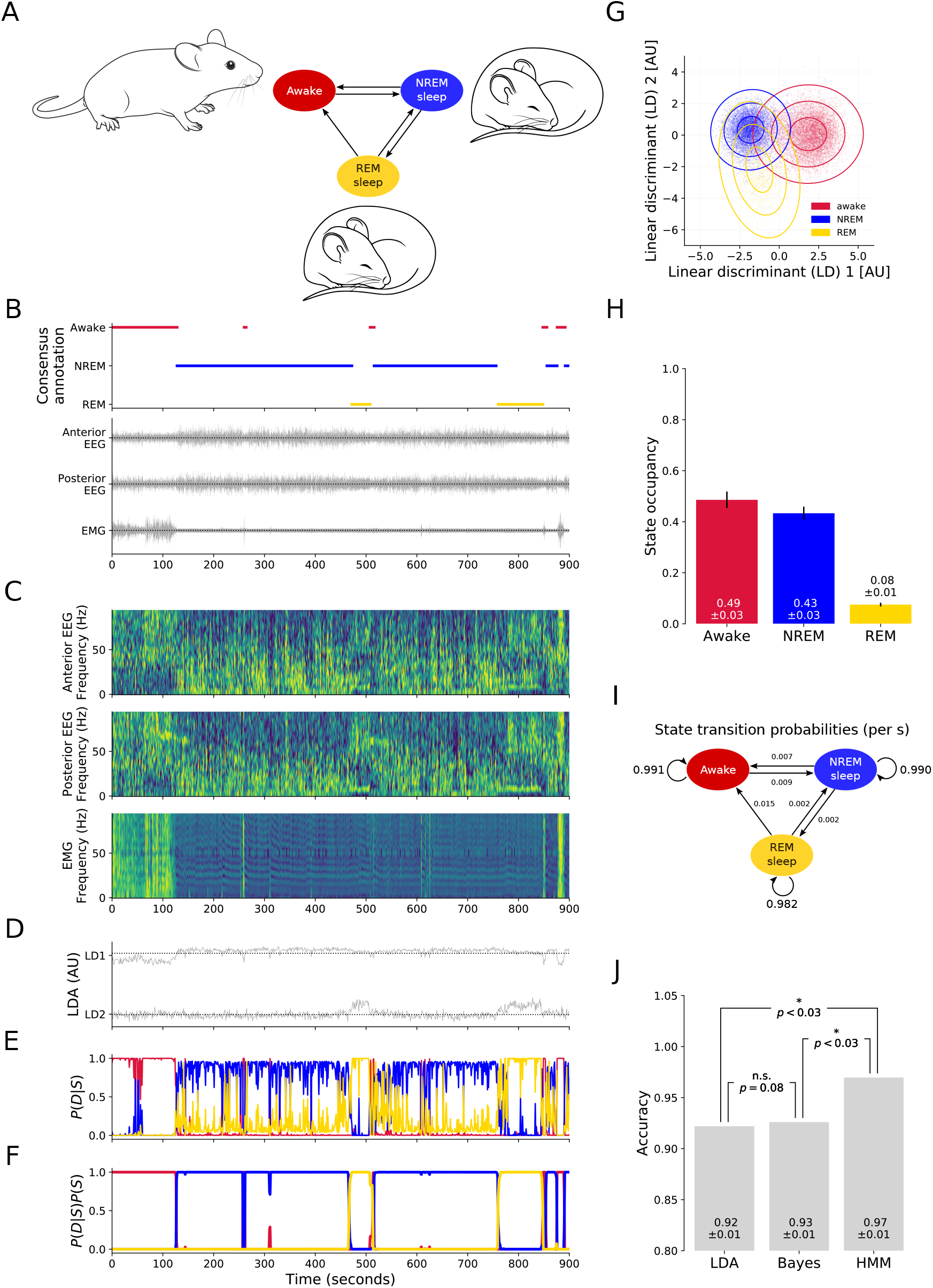
Establishing a probabilistic sleep stage classifier. (**A**) Continuous EEG and EMG recordings were made across a full sleep-wake cycle from freely behaving mice. (**B**) A fifteen-minute segment of the consensus of manual annotations by four independent experienced sleep researchers (top) and the corresponding anterior EEG, posterior EEG, and EMG recording. (**C**) Anterior EEG, posterior EEG and EMG multi-taper spectrograms. (**D**) Two-dimensional representation of the segment after targeted dimensionality reduction via LDA. Negative values in the first component (‘LD1’) and in the second component (‘LD2’) indicate the awake state; positive LD1 with negative LD2 indicates NREM; negative LD1 with positive LD2 indicates REM. (**E**) Probability of each state when fitting two-dimensional Gaussian distributions to the values in ‘D’. **(F)** Likelihood of each state given the probability of each state (as shown in ‘E’) and all possible state sequences, weighted by their likelihood given the state transition probabilities (as shown in ‘I’). (**G**) Distribution of values after dimensionality reduction by LDA. Each dot corresponds to a randomly chosen 1-second epoch. Colour indicates the state assigned in the manual consensus annotation. Lines indicate the standard deviations of multivariate Gaussian distributions, one for each state, fitted to all samples in the data set. (**H**) The state occupancy based on the time spent in each state across six 24-hour data sets, according to at least four manual annotations. (**I**) The corresponding state transition probabilities. (**J**) Accuracy of the LDA classifier, the naive Bayes classifier, and the HMM classifier (i.e. Somnotate). Accuracy was evaluated across six 24-hour data sets in a hold-one-out fashion. Error bars indicate standard deviation. P-values are derived from a Wilcoxon signed rank test with a Bonferroni-Holm correction for multiple comparisons.

The LDA classifier was found to achieve an accuracy of 92% ± 1% on the test data set (**Fig 1J, “LDA”**), which was comparable to previous work that used an LDA classifier to predict vigilance states from rodent experimental EEG data (89% ± 1% accuracy (22)). The Bayes classifier achieved an accuracy of 93% ± 1% on the test data set (**Fig 1J, “Bayes”**), which was consistent with previously reported values using a similar approach to predict vigilance states from rodent EEG data (94% ± 1% (23)). Finally, the HMM had an accuracy of 97% ± 1%, which was significantly more accurate than either the LDA or the Bayes classifiers, reducing the number of errors by more than half in both cases (**Fig 1J, “HMM”**). These analyses confirmed the benefit of incorporating contextual information. They also established automated feature extraction using LDA, combined with context-aware state annotation using a HMM, as a highly effective strategy for achieving very accurate automated sleep stage classification of experimental data. We refer to this improved classifier as ‘Somnotate’.

### Sleep stage annotation by Somnotate exceeds manual accuracy

Manual annotation continues to be the gold standard by which any automated annotation is measured. The performance of sleep stage classifiers is typically measured by computing their agreement with two independent manual annotations. Performance is evaluated as the average agreement of the automated annotation with each of the two manual annotations, and this average is then compared to the level of agreement between the two manual annotations. This subtle difference in how manual and automated annotations are compared can lead to systematic biases in favour of the automated annotation (see **Appendix S1**).

For these reasons, we were keen to compare manual and automated annotations to a majority-vote consensus derived from multiple independent manual annotations. We generated a data set of six 24-hour (i.e. 144 hours total) simultaneous recordings of anterior EEG, posterior EEG, and EMG in freely behaving mice (Materials and methods), which were independently scored by at least four experienced sleep researchers to annotate awake, NREM, and REM states. The sleep researchers had a median of 5 years’ experience in manual vigilance state annotation (minimum of 2 years’ experience), and had manually annotated a median of 1272 hours (minimum of 768 hours) of equivalent recordings (**Supplementary Table S1**). The recordings, individual manual annotations, and automated annotations are made freely available in standard formats at [WEBLINK].

To determine the accuracy of manual annotations by experienced sleep researchers, we compared individual manual annotations for any of the six 24-hour recordings to the consensus of the remaining three or more annotations for that data set. Somnotate was trained and tested in a hold-one-out fashion on the same data set, and its accuracy was determined by comparison to the consensus of the manual annotations. This revealed that the accuracy of Somnotate exceeded the accuracy of manual annotations by 13 experienced sleep researchers (**Fig 2A**). Out of a total of 25 manual annotations, 22 were less accurate than the automated annotation. Twelve out of the thirteen annotators had a lower average accuracy than the automated classifier on the same data sets. Thus Somnotate significantly exceeded human performance (p < 0.001, Wilcoxon signed rank). When we used Cohen’s kappa instead of accuracy as a measure of performance, we obtained identical results (**Supplementary Fig S7**).

**Fig 2.**
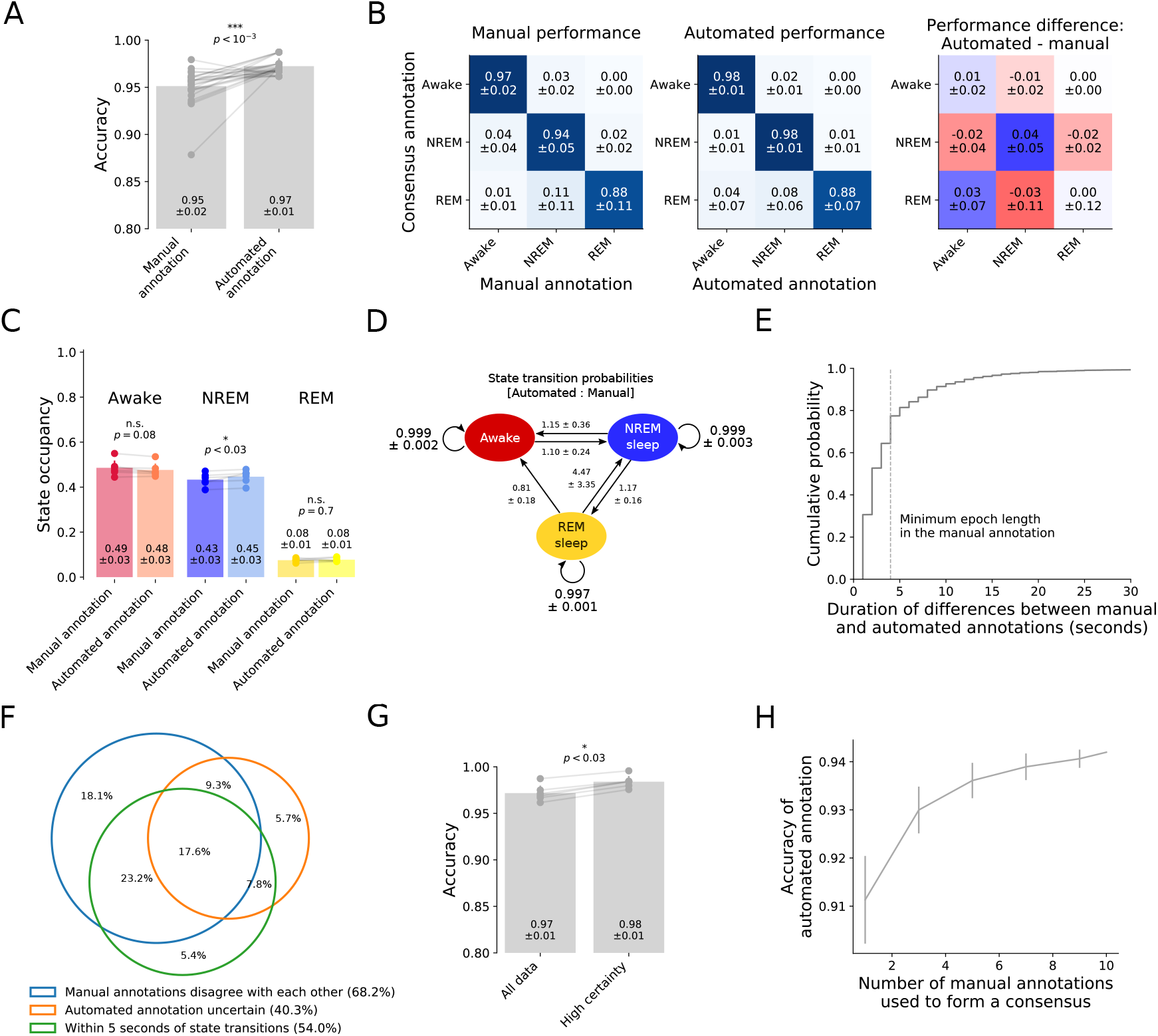
Automated sleep stage classification by Somnotate exceeds manual accuracy. (**A**) Somnotate was trained and tested, in a hold-one-out fashion, on six 24-hour data sets. Using a consensus annotation based on at least three manual annotations, the accuracy of the classifier was compared to the accuracy of individual manual annotations (n=25 manual annotations from 13 experienced sleep researchers). (**B**) The confusion matrix for individual manual annotations compared to the manual consensus (left), for the automated classifier compared to the manual consensus (middle), and the difference between these two confusion matrices (right). (**C**) Comparison of state occupancies between the automated and manual consensus annotations. (**D**) State transition probabilities in the automated annotation, normalised to the state transition probabilities in the manual consensus annotation. (**E**) Cumulative frequency plot shows the duration of the differences between the automated annotation and the manual consensus. Note that the manual annotation had a temporal resolution of 4 s (vertical dashed line), whereas the automated classification was performed at a time resolution of 1 s. (**F**) Venn-diagram of the time points at which the automated annotation and manual consensus differed. (**G**) Excluding samples where Somnotate is not certain improves accuracy. Classifier accuracy was compared between cases when all samples were included (‘All data’) and when 5.5% of samples were removed because the likelihood of the predicted state dropped below 0.995 (‘High certainty’). The plot indicates mean ± standard deviation and p-values are derived from a Wilcoxon signed rank test. (**H**) Somnotate was trained on six 24-hour data sets and then tested on a 12-hour data set, which had been independently annotated by ten experienced sleep researchers (as in Fig 1). The accuracy of the annotation by Somnotate was compared to consensus annotations generated from different numbers of manual annotations. Error bars indicate standard deviation. P-values are derived from a Wilcoxon signed rank test.

The difference between the confusion matrices for the manual and automated annotations indicated that the performance difference between manual and automated annotation was mainly driven by a more accurate annotation of NREM states (**Fig 2B**). Somnotate identified more state transitions than were typically present in manual annotations, in particular if these transitions involved NREM states. Cumulatively however, the differences between manual and automated state annotations resulted in minor differences in the overall state occupancy (**Fig 2C-D**). When partitioning the data by state according to the manual consensus or the automated annotation, there were no discernible differences between power spectra of the EEG activity (**Supplementary Fig S2**).

Approximately two thirds of the differences between Somnotate and the manual consensus annotations had a duration that was shorter than the temporal resolution of the manual annotations (i.e. shorter than 4 s; **Fig 2E**). Many of these differences could therefore be resolved if the data had been manually annotated at a higher temporal resolution, albeit at the expense of a greater investment of time. In other cases however, annotating a definitive state may not have been possible. For instance, the animal may have been transitioning from one state to another, resulting in ambiguous EEG and EMG waveforms that reflect an ‘intermediate’ state. Consistent with this scenario, 54% of differences between the consensus and the automated annotation occurred within 5 seconds of a state transition, where manual annotations also often disagreed with one another (**Fig 2F**). As Somnotate is a probabilistic classifier, it can automatically identify intermediate states as epochs with non-zero probabilities for more than one state (denoted by “Automated state annotations uncertain” in **Fig 2F**). Excluding these epochs reduces the differences between the consensus annotation and the automated annotation by a third (**Fig 2G**), and provides a convenient way to clean up the EEG data for further downstream analysis. As a final validation, we trained Somnotate on the six 24-hour test data sets, but then tested performance on the 12-hour data set that had been annotated by ten experienced sleep researchers. This revealed that the more manual annotations that were used to generate a consensus sequence of the test data, the more closely this manual consensus matched the automated annotation (Spearman’s rank correlation ρ = 0.80, p < 0.001; **Fig 2H**). In other words, as one increases the number of experienced annotators, the manual consensus converges on the automated annotation by Somnotate.

### Somnotate is highly robust

Machine learning algorithms can be uniquely sensitive to patterns in the data. This sensitivity is often desirable, but can also be problematic. We were therefore keen to examine Somnotate’s performance under conditions in which the training data contained errors, or where the test data reflected different experimental conditions, or where the data was acquired using a different experimental recording configuration.

First, to test sensitivity to errors in the training data, we evaluated Somnotate’s accuracy on six 24-hour EEG and EMG recordings annotated by at least four experienced sleep researchers in a hold-one-out fashion, while randomly permuting an increasing proportion of the consensus state annotations. This revealed that Somnotate is extremely robust to errors in the training data, as the accuracy on the test set only displayed a notable drop in performance when more than half of the training samples were misclassified (**Fig 3A-B**). Furthermore, classifier performance monotonically increased with increasing amounts of training data (**Supplementary Fig S3**), consistent with the idea that the classifier does not overfit the training data and, as a result, does not learn patterns that are present due to chance.

**Fig 3.**
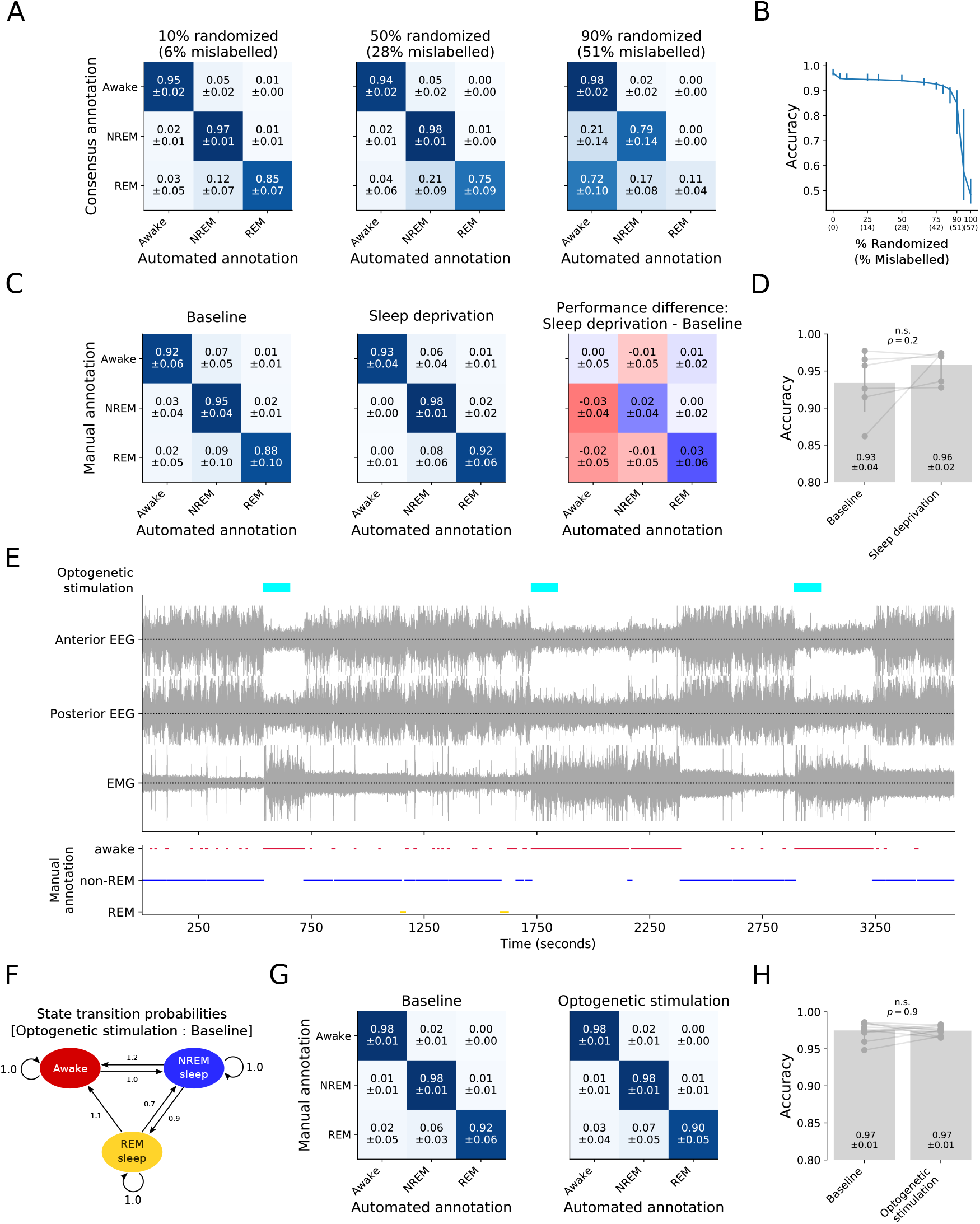
Somnotate is robust to errors in the training data, changes in the features of the data, and changes in the vigilance state transition probabilities. (**A**) Somnotate’s accuracy was evaluated on six 24-hour data sets, in a hold-one-out fashion, while permuting an increasing fraction of annotations in the training data. Confusion matrices show the results when permuting 10% of the training data annotations (resulting in 6% mislabelled time points; left), permuting 50% of the training data annotations (resulting in 28% mislabelled time points; middle), or permuting 90% of the training data annotations (resulting in 51% mislabelled time points; right). Values represent mean ± standard deviation. (**B**) Somnotate’s accuracy as a function of the percentage of permuted training data annotations. (**C**) The accuracy of Somnotate, pre-trained on 24h standard sleep-wake cycle datasets, was evaluated against a manual annotation of baseline control data (six 12-hour light cycle-only data sets), and compared to its accuracy on data from the same animals after undergoing a sleep deprivation protocol (six 12-hour light cycle-only data sets). Confusion matrices are shown for the baseline (left), following sleep deprivation (middle), and as the difference between these two confusion matrices (right). (**D**) Comparison of Somnotate’s overall accuracy on baseline data and data collected after sleep deprivation. P-value is derived from a Wilcoxon signed rank test. (**E**) Mice experienced experimentally-induced awakenings via optogenetic stimulation of ChR2-expressing inhibitory neurons in the lateral preoptic hypothalamus. (**F**) This optogenetic manipulation increased the probability that the animals transitioned from NREM sleep and REM sleep, to the awake state. (**G**) The accuracy of Somnotate, pre-trained on 24h standard sleep-wake cycle datasets, was evaluated against a manual annotation for eleven 24-hour data sets recorded during optogenetic stimulation, and for baseline recordings from the same animals on days when optogenetic stimulation was not performed. Confusion matrices are shown for the baseline recordings (left) and the recordings with optogenetic stimulation (right). (**H**) The accuracy of Somnotate’s annotations was near identical for both data sets and not significantly different (p > 0.9, Wilcoxon signed rank test).

Second, classifiers are susceptible to being fine-tuned to a standard training data set, such that performance levels can drop when faced with test data collected under different conditions, particularly when features used by the classifier are altered in a consistent manner. To assess robustness to changes in features, we tested Somnotate’s performance on data from a sleep deprivation experiment. Sleep deprivation is a common experimental manipulation that is known to change the EEG power spectrum after sleep onset, and EEG power values are primary features used by Somnotate. We evaluated the accuracy of our pre-trained classifier on six 3-hour data sets recorded after sleep onset in sleep-deprived mice, and compared this to the accuracy on matched ‘baseline’ (i.e. without sleep deprivation) data recorded from the same animals (**Fig 3C-D**). The annotation accuracy was found to be comparable and high across both the sleep deprivation and baseline conditions, consistent with Somnotate being robust to experimentally-induced changes in relevant features (**Fig 3D**).

Third, Somnotate uses contextual information in the form of prior probabilities of the different vigilance states. These probabilities depend on how much time animals spend in each state and how frequently they transition between states, both of which can change under experimental conditions. To assess the classifier’s robustness to variations in these prior probabilities, we trained and tested Somnotate in a hold-one-out fashion on the six 24-hour EEG and EMG recordings with high quality consensus annotations, as before. We then evaluated Somnotate’s accuracy on data sets from mice that had experienced repeated, experimentally-induced awakenings throughout the day. These awakenings were achieved via optogenetic stimulation of channelrhodopsin-2 (ChR2) expressing inhibitory neurons in the lateral preoptic hypothalamus (**Fig 3E**; as described in (42)), which affected the probabilities with which the animals transitioned between states (**Fig 3F**), but did not affect the overall time spent in each state (p > 0.15 for all states, Wilcoxon signed rank test, data not shown). Despite the changes in transition probabilities, there was no detectable change in Somnotate’s performance (**Fig 3G-H**).

Finally, although most electrophysiological signals show a dependence on vigilance state, some signals are thought to be more informative of certain states. For example, REM sleep is indicated by a high power in the theta frequency band of an EEG recording, which is typically more apparent for a posterior electrode than an anterior electrode (43). However, due to experimental requirements and technical limitations, it is not always possible to record the ‘ideal’ combination of signals for the inference of vigilance states. We were hence keen to assess Somnotate’s robustness to changes in recording configurations, and trained and tested Somnotate in a hold-one-out fashion on the six 24-hour recordings with high quality consensus annotations, while using only data from a single EEG, EMG, or LFP electrode. Our tests revealed that a single EEG signal was sufficient to infer the vigilance state with high accuracy (**Supplementary Fig S4**).

Taken together, these observations demonstrate the robustness of Somnotate to changes in the features used for inference, the state frequencies, and the state transition probabilities. Somnotate is therefore able to perform highly accurate sleep stage annotation under a variety of experimental conditions.

### Somnotate identifies failed state transitions

HMMs belong to the category of Bayes classifiers that compute the likelihood of each state, for every data sample (i.e. time point). This means that Somnotate is able to distinguish samples where it is certain in its prediction (i.e. where the likelihood of the predicted state is effectively one), and samples where it is uncertain (i.e. where the likelihood of the predicted state is less than one). For our data sets, the cumulative distribution of likelihood values indicated a change point at 0.995 (**Supplementary Fig S5**), with a minority of samples (5.5%) having a likelihood below this threshold. Notably, for samples where Somnotate was uncertain, almost half (44%) coincided with instances where manual annotations disagreed with one another. This suggested that the difficulty in predicting the vigilance state at these time points was not an artefact of the inference method, but a result of ambiguity within the signals. Human annotators often exclude such sections of the data from their analysis and, by analogy, the accuracy of our automated classifier further increased when these ambiguous samples were excluded **(Fig 2G)**. By reporting the certainty of its predictions, Somnotate greatly facilitates the identification of ambiguous samples that the experimentalist may wish to review or examine further. Somnotate also offers the chance to characterise such ambiguous samples in a principled way.

An ambiguous sample could result from measurement noise masking a true, unambiguous signal, or it could stem from a mixed signal reflecting an intermediate state. Four lines of evidence support the idea that ambiguous samples reflect an intermediate state and are not a measurement artefact. First, the majority of ambiguous samples (76%) occur around state transitions, which represent relatively rare events in the recordings (first two examples in **Fig 4A**). Second, for nearly all ambiguous samples, the probability mass was concentrated in two states, rather than being randomly distributed across all three states (**Fig 4B**). Third, the power spectra for ambiguous samples showed elements of the power spectra of the two most likely states (**Fig 4C**). For example, samples that Somnotate was uncertain whether to assign as awake or NREM sleep showed a high power in the δ frequency band, characteristic of NREM sleep, but also high power in the γ frequency band, which is typically an indicator of the awake state (**Fig 4C**). Fourth, the periods during which the classifier was uncertain tended to be much longer than the duration of a single sample and up to a maximum of 30 seconds, such that these intermediate states could not simply reflect the temporal resolution of the sampling (i.e. the result of a state transition occurring during a single one second sample).

**Fig 4.**
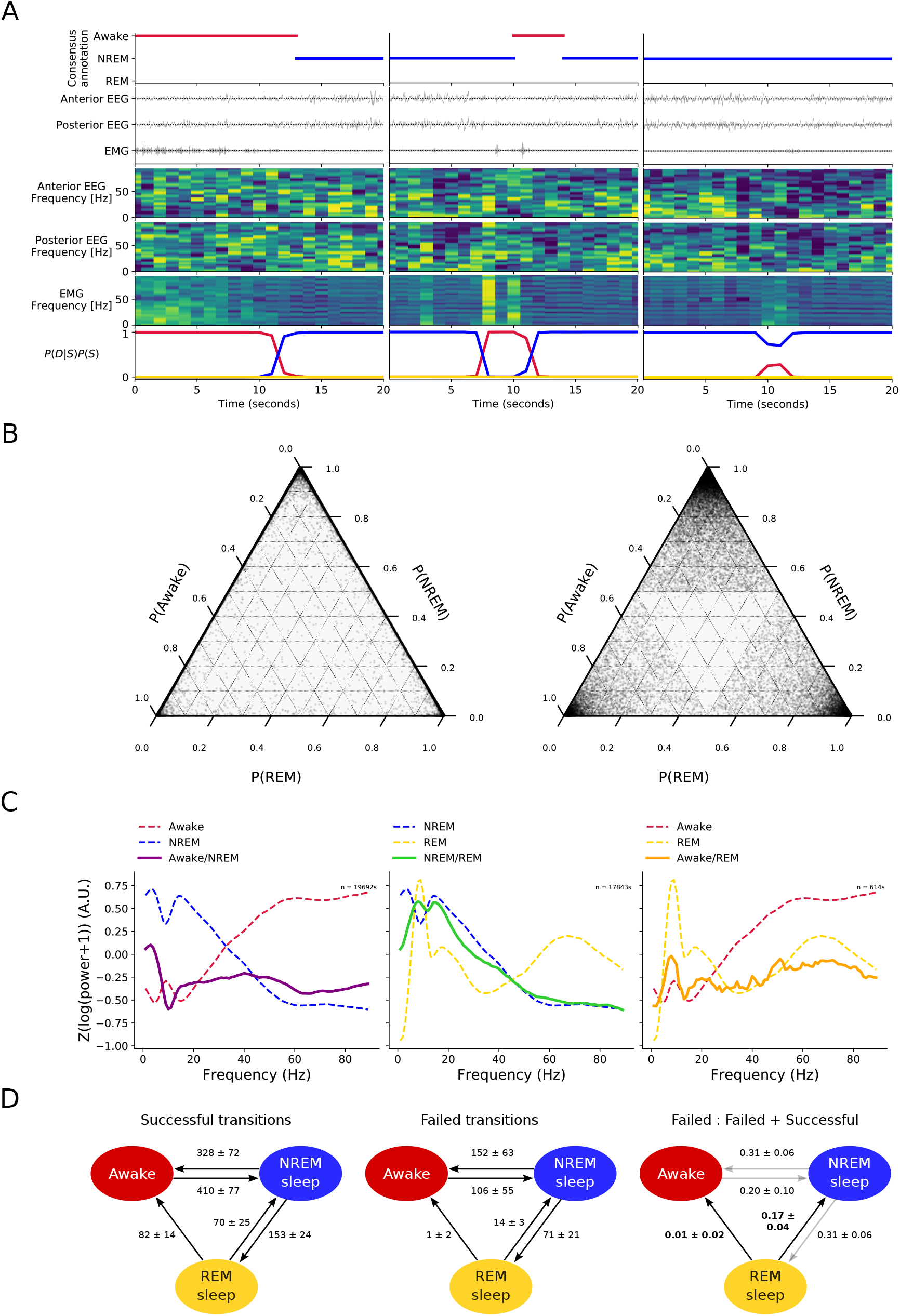
Somnotate identifies intermediate states associated with successful and failed vigilance state transitions. (**A**) Three examples of intermediate states identified by Somnotate, in which the probability of the most likely state dropped below 0.995. In each case, the consensus annotation, input signals, power spectra and likelihood of each state assigned by Somnotate, are shown. The first example (left) shows a successful state transition from awake to NREM sleep. Just before the transition, Somnotate identifies time points with intermediate states in which the probability of being awake has decreased and NREM sleep has increased. The second example (middle) shows a brief state transition from NREM sleep to awake, and then back to NREM sleep, which includes time points with intermediate states. The third example (right) shows a failed transition from NREM sleep to awake, which includes a series of time points with intermediate states in which there is a partial decrease in the probability of NREM sleep and partial increase in the probability of being awake. (**B**) Ternary plot of the state probabilities assigned to each time point with an intermediate state in six 24-hour data sets (left). In the vast majority of cases, the probability mass was concentrated in one or two states. This was different to a theoretical distribution in which the probability mass outside the most likely state was randomly assigned to the other two states (right). (**C**) Power spectra extracted for time points with intermediate states (solid lines). For reference, the power spectra for the “pure” states are also shown (dashed lines). (**D**) Relative frequencies of successful state transitions (per day; left), failed state transitions (middle) and the ratio between these (right). Values indicate mean ± standard deviation. The failure rates of REM transitions were statistically significantly different from one another (p < 0.001; χ^2^ contingency test), and hence indicated with black arrows; the failure rates of NREM transitions were not significantly different (p > 0.05; χ^2^ contingency test), and hence indicated by grey arrows.

To demonstrate the opportunities afforded by Somnotate’s identification of intermediate states, we focused upon the 24% of ambiguous samples that were associated with an incomplete state transition (such as the third example in **Fig 4A**) and referred to these as ‘failed transitions’, to distinguish them from successful state transitions. We computed the frequencies of these different transition types and expressed the failed transitions as a proportion of all transitions (**Fig 4D**). This revealed that the probability of failing to transition was not random. The overall probability of failed transitions was higher when moving out of NREM sleep, than when moving out of REM sleep (p < 0.001, χ^2^ contingency test). Furthermore, whereas the two state transitions out of NREM sleep exhibited a similar probability of failing (p = 0.98, χ^2^ contingency test), the state transitions out of REM sleep differed, with a transition from REM-to-NREM showing a higher probability of failing than a transition from REM-to-awake (17% versus 1%; p < 0.001, χ^2^ contingency test). The same pattern of failed transitions remained when a more conservative threshold criterion was adopted such as a state probability below 0.95 (data not shown). These observations may explain why animals often appear to enter a brief awake state between REM sleep and NREM sleep (33–36), as transitions from REM to awake, and from awake to NREM, fail much more rarely than transitions from REM to NREM.

Finally, we investigated if the state transition failure rates were sensitive to an animal’s recent sleep-wake history. The recovery after sleep deprivation is known to be dominated by long periods of NREM and REM sleep **(Fig 5A** & (2,33)). We asked whether this change in state occupancy is also associated with a change in the failure rate of state transitions. Indeed, we observed an increase in the failure rate of transitions out of NREM, as well as an increase in the failure rate of REM-to-NREM transitions, increasing the likelihood of remaining in either sleep state for a longer period of time (p < 0.05, Mann-Whitney rank test, **Fig 5B**). In contrast, there was no difference in the failure rate for awake-to-NREM transitions. Together, these changes in the failure rate of the different state transitions are consistent with the observed vigilance state occupancy following sleep deprivation. More generally, these analyses confirm the additional opportunities afforded by Somnotate’s ability to identify intermediate states and to provide a more complete and richer characterisation of sleep-wake dynamics.

**Fig 5.**
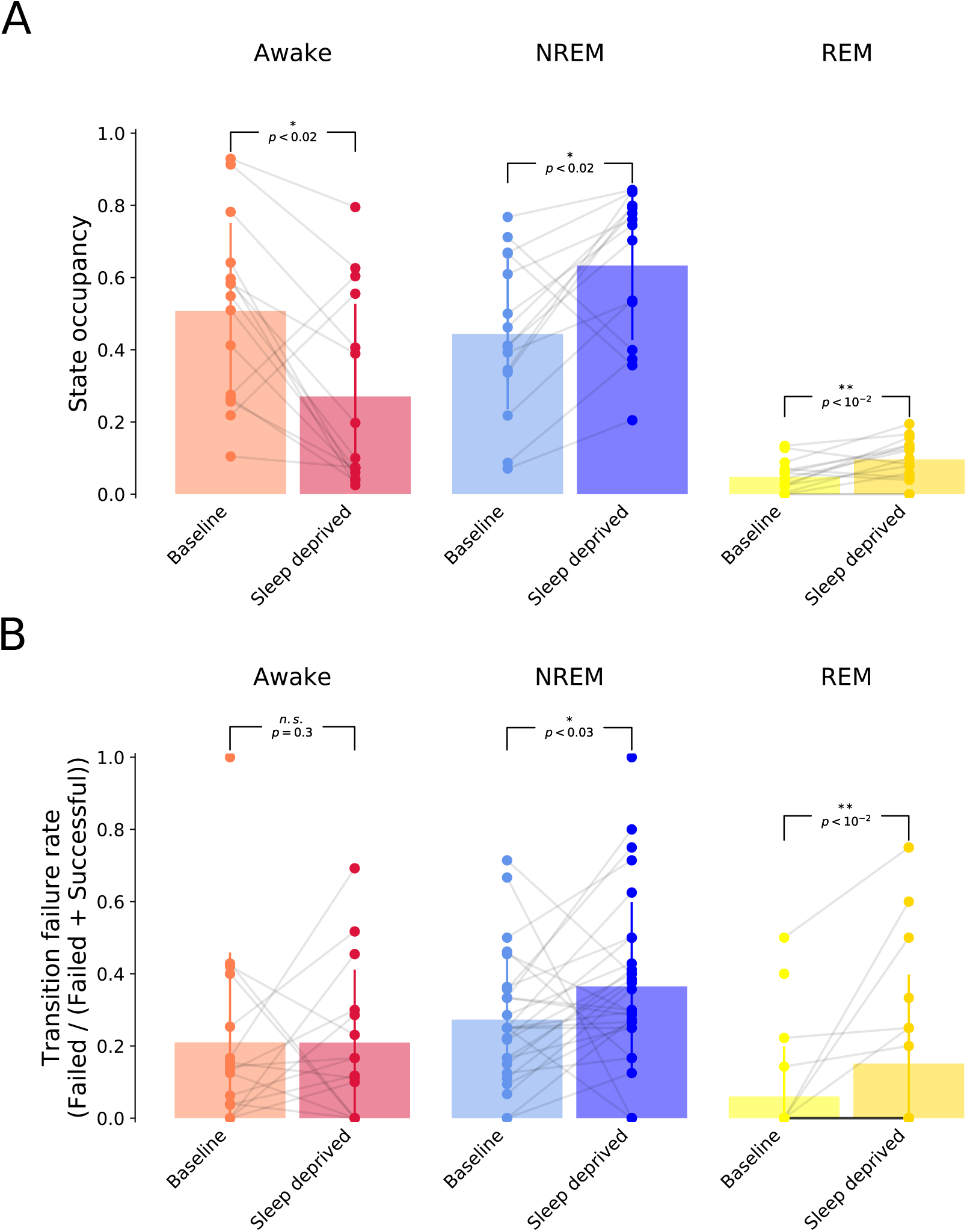
**Vigilance state transition failure rates depend upon sleep-wake history. (A**) State occupancy during the first two hours of recovery from a period of sleep deprivation versus baseline recordings performed in the same animals over an equivalent period but not following sleep deprivation. (**B**) Failure rates for transitions during the awake state (i.e. awake-to-NREM transitions; left), transitions during NREM (NREM-to-awake and NREM-to-REM transitions; middle), and transitions during REM (REM-to-awake and REM-to-NREM transitions; right). P-values are derived from a Wilcoxon signed rank tests with a Bonferroni-Holm correction for multiple comparisons.

## Discussion

We present Somnotate - a purpose-built probabilistic sleep stage classifier that quantifies the certainty of its annotations. This enables the experimenter to assign vigilance states (awake, REM sleep, and NREM sleep) with high accuracy, whilst also identifying ambiguous epochs that represent intermediate states. Somnotate combines optimal feature extraction by linear discriminant analysis (LDA), with state-dependent contextual information derived via a hidden Markov model (HMM). Our benchmarking tests demonstrate that this approach optimises the use of contextual information, outperforming other classifiers that are compatible with data collected under experimental conditions. Furthermore, through systematic comparisons against expert manual annotations, we demonstrate that Somnotate surpasses the accuracy achieved by experienced sleep researchers. Somnotate is shown to be robust to errors in the training data, able to operate across different types of experimental manipulations, and compatible with different electrophysiological signals.

Somnotate’s ability to quantify the certainty of its predictions creates new opportunities for the researcher, by systematically identifying epochs in which the vigilance state is ambiguous. With its dual output at every time point – a prediction and a certainty for each vigilance state – Somnotate offers users the chance to rapidly identify and confirm the classifier’s performance in a targeted manner. Furthermore, this feature affords new analysis opportunities that are only feasible with an automated approach. Whilst the gold standard for sleep stage classification remains human experts, there is an element of subjectivity to all manual annotations that makes areas of investigation difficult. For example, EEG signals often show signatures of multiple states, particularly around state transitions (30,44–46), where most disagreements between manual annotations occur. And although humans are very good at determining the most likely state at a given time point, they struggle to quantify intermediate states.

To demonstrate the opportunities that result from this feature, we used Somnotate to identify intermediate states associated with incomplete state transitions, which we define as failed transitions. Interestingly, the probability of failed transitions was highly non-random and depended upon the type of transition, with REM-to-awake transitions showing relatively low failure rates, but REM-to-NREM transitions showing high failures. This differential failure rate may explain the preponderance of brief awake periods between REM and NREM sleep, as it may be easier for the underlying neuronal networks to transition from REM to awake, and then to NREM, rather than to transition directly from REM to NREM (33–36). In addition, we revealed that the animal’s recent sleep-wake history affects the probability of observing different types of failed transitions. More specifically, recovery sleep following a period of sleep deprivation exhibited an increase in the probability of failed transitions from NREM to REM, and from NREM to awake. These observations suggest that the underlying neuronal networks find it more difficult to transition out of the NREM state following sleep deprivation, which is consistent with the well-established finding that sleep deprivation leads to longer and more intense periods of NREM sleep (2,33). These applications highlight the power of Somnotate’s ability to determine intermediate states and reveal new opportunities to investigate the neurophysiological mechanisms of state transitions.

Despite previous efforts to develop algorithms for sleep stage classification, many sleep researchers continue to manually score their data. We believe that several barriers have prevented widespread adoption of automated solutions and our findings allow us to evaluate Somnotate in terms of these potential stumbling blocks. A first barrier to the widespread adoption of automated solutions is the issue of accessibility. Commercially available software for automated sleep stage classification can be expensive and details of their operation are often not made available to the user (26,47). We provide an open source implementation of our algorithm written in Python. The code comes with extensive documentation including detailed installation instructions and a comprehensive tutorial. The modules of the code base can be integrated into an existing workflow. Alternatively, we also provide a fully-fledged pipeline as a standalone command line application.

In terms of performance levels, reports of human-like performance by automated methods may fall short in practice. Our findings suggest that the choice of performance metric may have contributed to this, as we show that inter-rater agreement can be an imprecise measure of annotation accuracy and is typically used in a manner that favours automated annotation. To improve upon this standard, we evaluated the quality of automated annotations against the consensus of at least three manual annotations. An annotation based on the consensus by majority vote will be more accurate than any individual annotation, whenever manual errors show some degree of independence from one another (48,49). Using this improved assessment of performance, we showed that Somnotate matched the consensus more closely than any individual manual annotation. What was particularly encouraging was that the more manual annotations that were used to generate the consensus sequence, the more closely this consensus matched the automated annotation by Somnotate.

Human experts are aware that they continually use contextual information whilst interrogating time series data, relating information at a time point of interest, with information that they infer over longer timescales. This is often overlooked in automated classifiers, although a subset have used algorithms that incorporate contextual information, including those that have used HMMs (7,8) and recurrent neural networks (11,12,14–17). The inherent flexibility of recurrent neural networks can pose problems, as they require large amounts of labelled data to train, are prone to overfitting, and adapting their architecture to different inputs can be non-trivial. In a research setting, where changes to the experimental setup can be frequent and generating large, well-curated training data sets is often impractical, recurrent neural networks can be a suboptimal choice. As HMMs are easier to optimise by non-experts, place significantly less demands on training data, and are thus more compatible with the constraints of experimental settings, we concentrated our efforts on improving the state-of-the-art for HMM-based inference of vigilance states. Somnotate represents an advance upon previous work on related classifiers, by first using LDA to automatically extract features that carry the maximum amount of linearly decodable information about vigilance states, and then incorporating state-dependent contextual information through the application of a HMM.

Another concern with automated approaches is how annotation is affected by differences in the training data or the features used for classification. We found that Somnotate is remarkably robust to errors in the training data, with test performance only dropping significantly when more than half of the training samples had been deliberately misclassified. Furthermore, automated scoring methods can exhibit overfitting to standard or control data sets, which means that their performance is diminished in other settings when the probabilities of key features vary (14,17,19,50,51). This was not the case with Somnotate. We saw no drop in performance when Somnotate annotated data collected under different experimental conditions in which the features used for inference, the state transition probabilities, and/or the state frequencies varied. Somnotate’s high performance was maintained on data from sleep-deprived mice in which the EEG spectrogram is significantly altered, and on data from optogenetic manipulation experiments in which the state transition probabilities are changed. These applications establish that Somnotate is well-suited to perform accurate sleep stage annotation under a variety of experimental conditions.

In terms of adaptability, the use of targeted feature extraction via LDA means that Somnotate is agnostic with respect to the exact nature of the input signal, and our data suggests that Somnotate retains a high accuracy with different EEG recording configurations and even other data sources such as local field potential (LFP) recordings. In principle, Somnotate could be applied to any high frequency time series data that contains information about an animal’s vigilance state, and hence we plan to expand this approach to other types of signals, such as surface EEG, actigraphy, or respiratory activity (24,51–53). While some automated approaches depend on a specific input signal (25), others are also adaptable in the sense that they can be recalibrated to changes in the experimental setup (10,20,22–24,32). However, as these automated approaches tend to have several free parameters, adaptation to a different setup can be time-consuming, with uncertain returns. As there are no free parameters in Somnotate other than the desired time resolution of the state prediction, re-training requires no optimisation, and is straightforward and fast. Training Somnotate takes approximately one second per 24 hours of data on a standard desktop computer.

In summary, Somnotate’s development as a probabilistic sleep stage classifier affords new biological insights through the identification and characterization of intermediate states in polysomnography, whilst simultaneously setting new standards in terms of performance, robustness, ease of use, and accessibility.

## Materials and methods

### Animal husbandry and sleep deprivation

All experiments were performed on adult male C57BL/6 wild-type mice, which were bred, housed and used in accordance with the UK Animals (Scientific Procedures) Act (1986). Animals were maintained under a 12-hour:12-hour light-dark (LD) cycle. For the subset of animals that underwent a sleep deprivation (SD) protocol, each animal was pre-exposed to novel objects to encourage exploratory behaviour. The SD protocol then consisted of delivering novel objects for the first three (Fig 3) or for the first six hours (Fig 5) of the light cycle, under the continuous observation of an experimenter. Once an animal had stopped exploring an object, a new object was presented.

### Surgical procedures and electrode configuration

For chronic electroencephalogram (EEG) and electromyogram (EMG) recordings, custom-made headstages were constructed by connecting three stainless steel screw electrodes (Fine Science Tools), and two stainless steel wires, to an 8-pin surface mount connector (8415-SM, Pinnacle Technology Inc., Kansas). For LFP recordings, a 16-channel silicon probe (NeuroNexus Technologies Inc., Ann Arbor, MI, USA; model: A1×16-3mm-100-703-Z16) with a spacing of 100 μmm between individual channels was used. Device implantation was performed using stereotactic surgery, aseptic technique, isoflurane anaesthesia (3-5% for induction and 1-2% for maintenance) and constant body temperature monitoring. Analgesia was provided at the beginning of surgery and during recovery (buprenorphine and meloxicam). A craniotomy was performed over the right frontal cortex (AP +2 mm, ML +2 mm from Bregma), right occipital cortex (AP +3.5 mm, ML +2.5 mm from Bregma), and the cerebellum (-1.5 mm posterior from Lambda, ML 0). A subset of animals were further implanted with a bipolar concentric electrode (PlasticsOne Inc., Roanoke, VA, USA) in the right primary motor cortex, anterior to the frontal EEG screw. To accommodate this additional implant, the frontal EEG screw was typically implanted 0.2-1.6 mm posterior to the target coordinates. For EEG recordings, a screw was fixed over both the right frontal and occipital cortex. For LFP and multi-unit activity recording in a subset of animals, a 16-channel silicon probe was implanted into primary motor cortex (+1.1 mm AP (anterior), -1.75 mm ML (left), tilt -15° (left)) under microscopic control, as reported previously (54). EEG and LFP signals were referenced to a cerebellum screw. For EMG recordings, wire electrodes were inserted into the left and right neck muscles, and one signal acted as reference to the other. All implants were secured using a non-transparent dental cement (SuperBond from Prestige Dental Products Ltd, Bradford, UK). Animals were allowed to recover for at least 1 week before recordings.

### In vivo data acquisition

Animals were moved to a recording chamber and housed individually in a Plexiglas cage (20.3 x 32 x 35 cm). Recordings were performed using a 128-channel Neurophysiology Recording System (Tucker-Davis Technologies Inc., Alachua, FL, USA), acquired using the electrophysiological recording software, Synapse (Tucker-Davis Technologies Inc., Alachua, FL, USA), and stored locally for offline analysis. EEG, EMG, and LFP signals were continuously recorded, filtered between 0.1–100 Hz, and stored at a sampling rate of 305 Hz. EEG, EMG and LFP signals were resampled at 256 Hz using custom code in MATLAB (MathWorks, v2017a), and converted into the European Data Format. The first and/or last 30 seconds of recordings could contain missing values as this corresponded to the period when the electrodes were being connected/disconnected from the recording system. These epochs were excluded from all subsequent analyses.

### Optogenetic stimulation

We employed a protocol previously described in detail in (55) and (42). Briefly, channelrhodopsin-2 (ChR2) was expressed in glutamate decarboxylase 2 expressing (GAD2^+^) interneurons by injection of an adenovirus construct (UNC vector core, AAV5-EF1a-DIO-ChR2-eYFP) into the lateral preoptic area (LPO) of the hypothalamus in adult Gad2-IRES-Cre mice (Jackson Laboratory 019022; B6N.Cg-Gad2^tm2(cre)Zjh^/J). For optical stimulation, either an optic fiber (400 μmm diameter, Doric Lenses Inc, Quebec) or a custom made optrode, consisting of an optic fiber glued with tungsten wires, was inserted to 0.2 mm above the virus injection site. All electrophysiological recordings were made 4 to 7 weeks post virus injection. To optogenetically stimulate GAD2^+^ neurons, we applied 10 ms pulses of light from a blue LED (470 nm, 10.8 – 13.2mW at fiber tip) at various frequencies (20, 10, 5, 2 or 1 Hz), for a duration of 2 minutes, every 20 ± 2 minutes. On baseline days, no optogenetic stimulation was provided.

### Manual vigilance state annotation

Manual annotation of vigilance states was performed offline, based on 4 s epochs using SleepSign software (Kissei Comtec). The anterior EEG channel, the posterior EEG channel, and the EMG channel were displayed on-screen simultaneously and visually inspected for vigilance state scoring. Three vigilance states were identified, as is typical in laboratory rodent studies. Waking was defined by a low-voltage, high-frequency EEG signal, with a high level or phasic EMG activity. During active, exploratory waking, a transient increase in theta-activity (5-10 Hz) was typically observed in the occipital derivation, overlying the hippocampus. NREM sleep was defined by an overall higher amplitude signal, dominated by slow waves (<4 Hz) and spindle oscillations (10-15 Hz) that were especially prominent in the anterior EEG channel, while the EMG signal was typically low. REM sleep was characterised by low-voltage, high-frequency EEG, dominated by theta activity especially in the posterior EEG channel, with a low level of EMG activity.

### Data pre-processing for automated annotation

We first computed the spectrograms of the anterior EEG, the posterior EEG, and the EMG traces. To reduce sensitivity to noise present in electrophysiological recordings, we used a multitaper approach, as this results in more robust estimates of the power than the more conventional Baum-Welch algorithm. Specifically, we used the implementation in the lspopt python library (1 second long segments with no overlap, other parameters at default values). We then discarded parts of the power spectrum that are strongly influenced by signals not related to changes in vigilance states. We discarded signals in the 0-0.5 Hz frequency range in the EEG and EMG recordings, as these are dominated by drift due to animal locomotion. Furthermore, we discarded signals between 45-55 Hz and above 90 Hz, as these were strongly affected by 50 Hz electrical noise. We then applied a log(x+1) transformation to map the heavy-tailed distribution of power values to a distribution that is more normally distributed. The normal distribution is the maximum entropy distribution for continuous distributions on unbounded domains, and as such, samples are maximally far apart from one another (compared to other distributions with the same variance). This facilitates downstream classification into separable groups. The re-mapped power values were then normalised by converting them to Z-scores (mean subtraction followed by rescaling to unit variance). Normalisation ensures that all frequencies are weighted equally in the downstream feature extraction. Finally, the normalised spectrograms were concatenated, resulting in a high-dimensional signal.

### Automated feature extraction

Features for downstream classification were then extracted from the concatenated spectrograms in a targeted manner using linear discriminant analysis (LDA (57)), as implemented in the scikit-learn python library (58). LDA determines a linear projection of high dimensional data to a low dimensional representation, such that samples belonging to different classes are optimally linearly separated in the low dimensional space. Thus, information in the signal about the different classes is preserved, while non-informative components of the signal are discarded. This has two further effects. Firstly, training of any classifier is accelerated, which implicitly or explicitly fits a joint probability distribution to the components of the training data. The number of samples required to accurately fit a joint probability distribution increases exponentially with the number of dimensions. As the dimensionality of the data is reduced, fewer samples are required to escape the under-sampled regime and accurately determine the shape of the data distribution. This is enhanced by the fact that the components of the LDA are largely independent of one another – unlike the original signal, in which many frequencies are highly correlated with each other. Secondly, as much of the original signal is effectively discarded, artefacts that contaminate the signal are also removed.

### Automated vigilance state annotation

Given three target states (awake, NREM sleep, and REM sleep), dimensionality reduction with LDA results in two-dimensional signals. These two-dimensional signals together with the corresponding manual annotations were used to train a HMM in a supervised fashion, with multivariate Gaussian state emissions using the python library pomegranate all optional parameters at default values (59). If the annotations were not based on the consensus of multiple manual annotations, mislabelled samples in the training data resulted in non-zero probabilities for disallowed state transitions, specifically awake-to-REM transitions. These were pruned by removing all state transitions with probability below 0.0001 per second. The accuracy of the trained LDA and HMM models were ascertained by applying the models to held out test data. For each sample, the probability of each state was computed using the Baum-Welch algorithm, and the most likely state sequence was determined using the Viterbi algorithm. Unless specified otherwise, training and testing occurred in a hold-one-out fashion.

### Recording artefacts

Samples containing artefacts associated with the animal’s gross body movements were identified during manual annotations, but were still included in the analysis of vigilance states and in the data used to train Somnotate. Such artefacts represented 1.0% ± 1.0% of the consensus manual annotations (mean ± standard deviation; 3.8% ± 2.8% in the individual manual annotations) and did not influence the automated feature extraction by LDA, so did not impact the quality of the automated annotations. However, such artefacts could affect downstream analyses in future applications, such as spectral analysis of the recorded signals. For this reason, Somnotate includes two features to facilitate the detection and removal of artefacts. First, Somnotate detects and demarcates gross movements that generate voltage deflections outside of the dynamic range of the recording system (with an optional padding to also remove voltage deflections preceding and following such events), so that they are not included in downstream analyses. Second, Somnotate has the option to present samples to the user where the classifier was uncertain about state assignment. Intervals consisting of consecutive samples in which the probability of the inferred state is below one are scored according to the sum of the residual probabilities (i.e. one minus the probability of the inferred state) and presented to the user in descending order. Movement artefacts associated with prolonged voltage deflections, or that strongly affect the spectral features identified by LDA, result in a high score and can be excluded by the user.

## Author contributions

PJNB designed and wrote the Somnotate software. HA tested and provided feedback on Somnotate during its early development. PJNB and CJA designed the validation experiments and PJNB carried out the analysis. HA, LBK, CBD, and TY performed the in vivo recordings. LBK organised the manual annotation effort. HA, LBK, CBD, ASF, SF, MCCG, YGH, MCK, LEK, LEM, LM, LT, CWT, TY, and VVV contributed manual annotations. RGF, VVV and CJA supervised the work. PJNB and CJA wrote the manuscript, with input from all authors.

## Acknowledgements

We would like to thank the Akerman lab for advice and comments. The research leading to these results received funding from the European Research Council under the European Community’s Seventh Framework Programme FP7/2007-2013, ERC Grant Agreement 617670 and also Medical Research Council (UK) project MR/N026039/1 and MR/S01134X/1, and Wellcome Trust 106174/Z/14/Z. HA was funded by a Sir Henry Wellcome Postdoctoral Fellowship and St John’s College Junior Research Fellowship; LBK by a Wellcome Trust PhD studentship (203971/Z/16/Z), Hertford College Senior Scholarship, Goodger and Schorstein Research Scholarships in Medical Sciences; CBD by a Wellcome Trust PhD studentship (109059/Z/15/Z) and Clarendon Fund Scholarship; MCCG by a BBSRC DTP grant (BB/J014427/1) and Clarendon Scholarship; YGH by a Medical Research Council/Stroke Association Clinical Research Training Fellowship; MCK by a Berrow Foundation Lord Florey Scholarship, plus Goodger and Schorstein Research Scholarship in Medical Sciences; LEM by a Biotechnology and Biological Sciences Research Council Industrial CASE Grant with Eli Lilly & Company Ltd (BB/K011847/1), a Novo Nordisk Postdoctoral Fellowship and a Sir Paul Nurse Junior Research Fellowship at Linacre College; LM by an Action on Hearing Loss studentship; CWT by a BBSRC DTP Grant (BB/M011224/1); TY by a Uehara Memorial Foundation Overseas Postdoctoral Fellowship and a Naito Grant for Studying Overseas.

## Supplementary Information

**Supplementary Table S1.** **Manual annotations were performed by experienced sleep researchers.** All authors who provided manual annotations reported their task-relevant experience in years, plus the approximate number of hours that they had previously manually annotated.

| Annotator | Experience (Years) | Total data annotated (Hours) |
| --- | --- | --- |
| VVV | 22 | 10000 |
| TY | 6 | 3840 |
| LEM | 7 | 2400 |
| LT | 5 | 2400 |
| CBD | 6 | 1608 |
| HA | 3 | 1368 |
| CWT | 3 | 1344 |
| MCCG | 6 | 1200 |
| YGH | 5 | 1200 |
| MCK | 4 | 1200 |
| LBK | 5 | 1200 |
| SJF | 2 | 960 |
| ASF | 2 | 840 |
| LM | 5 | 768 |
| Mean | 5.79 | 2166 |
| Median | 5 | 1272 |
| Minimum | 2 | 768 |

**Supplementary Fig S2.**
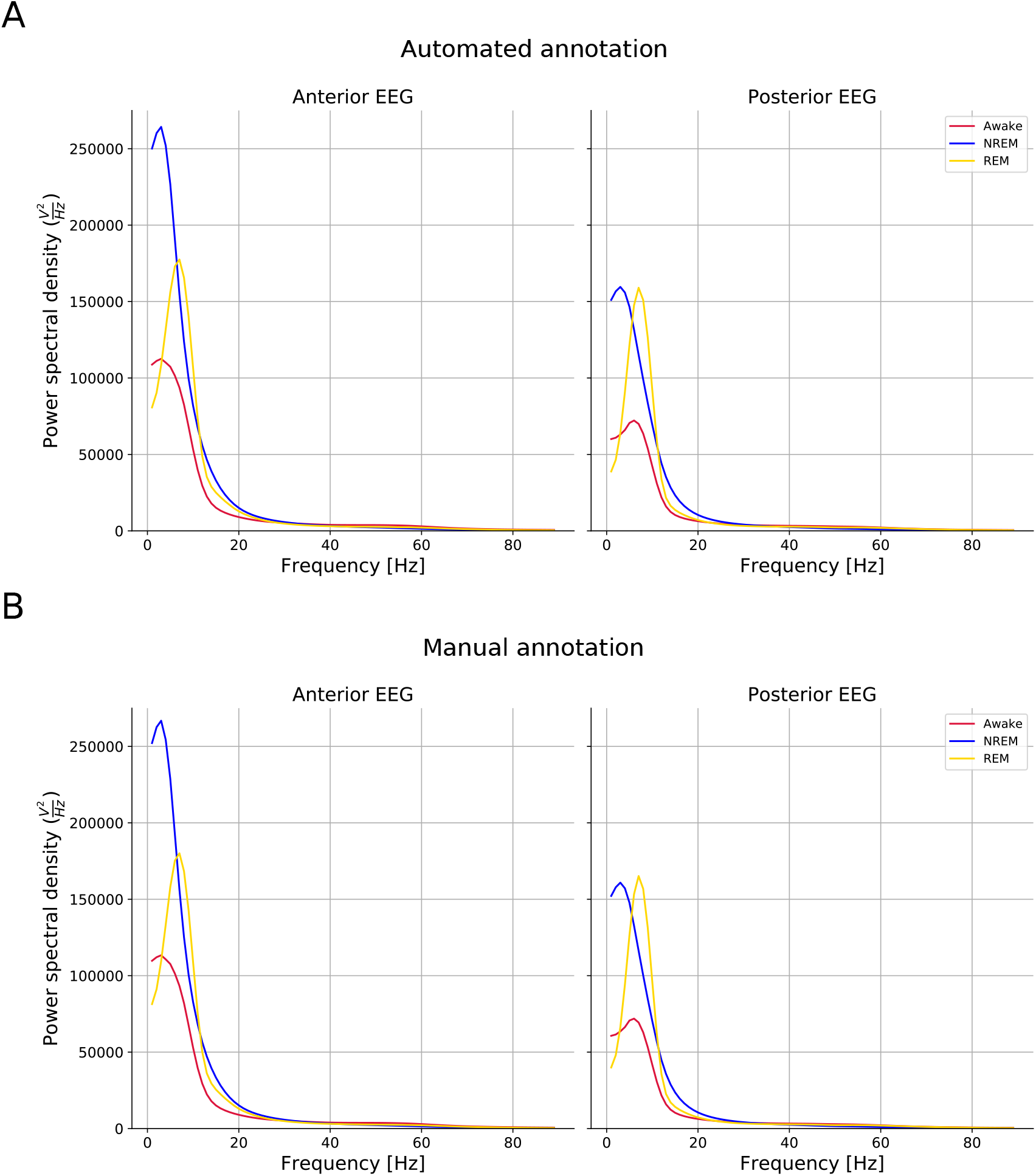
EEG power spectra by state according to Somnotate (A) or manual consensus annotations (B). Somnotate was trained and tested, in a hold-one-out fashion, on six 24-hour data sets. Spectrograms were computed for the anterior and posterior EEG and partitioned according to the predicted state. The process was repeated using the manual consensus annotations. The lines indicate the median EEG power.

**Supplementary Fig S3.**
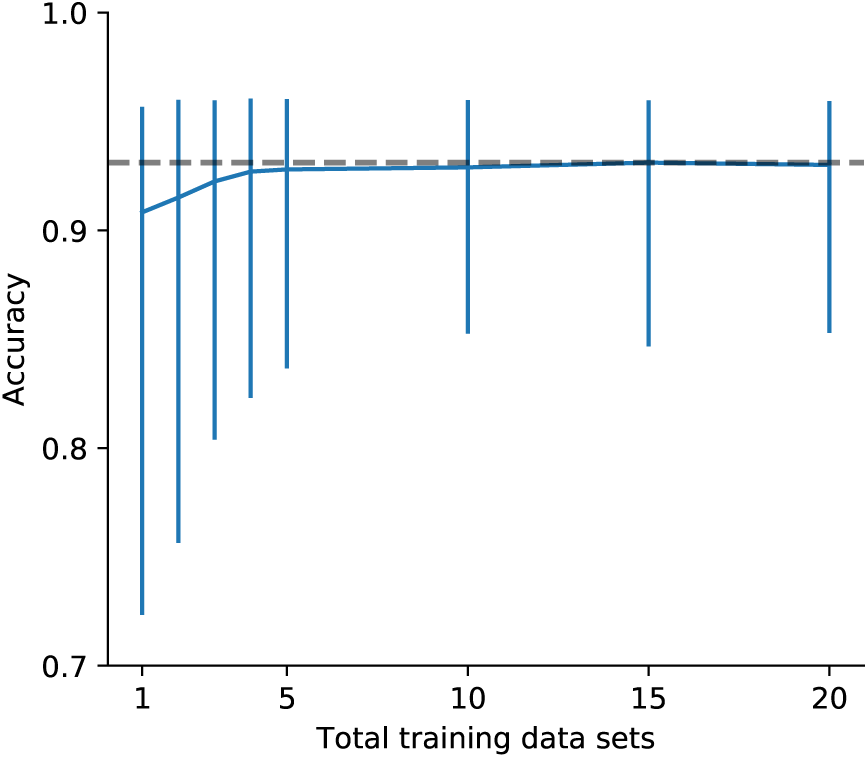
Somnotate’s performance as a function of the amount of training data. Somnotate was trained and tested, in a hold-one-out fashion, using different numbers of 24-hour data sets. The maximum amount of training data available was twenty 24-hour EEG and EMG recordings under baseline conditions. The line indicates the median. Error bars demarcate the 5th and 95th percentile.

**Supplementary Fig S4.**
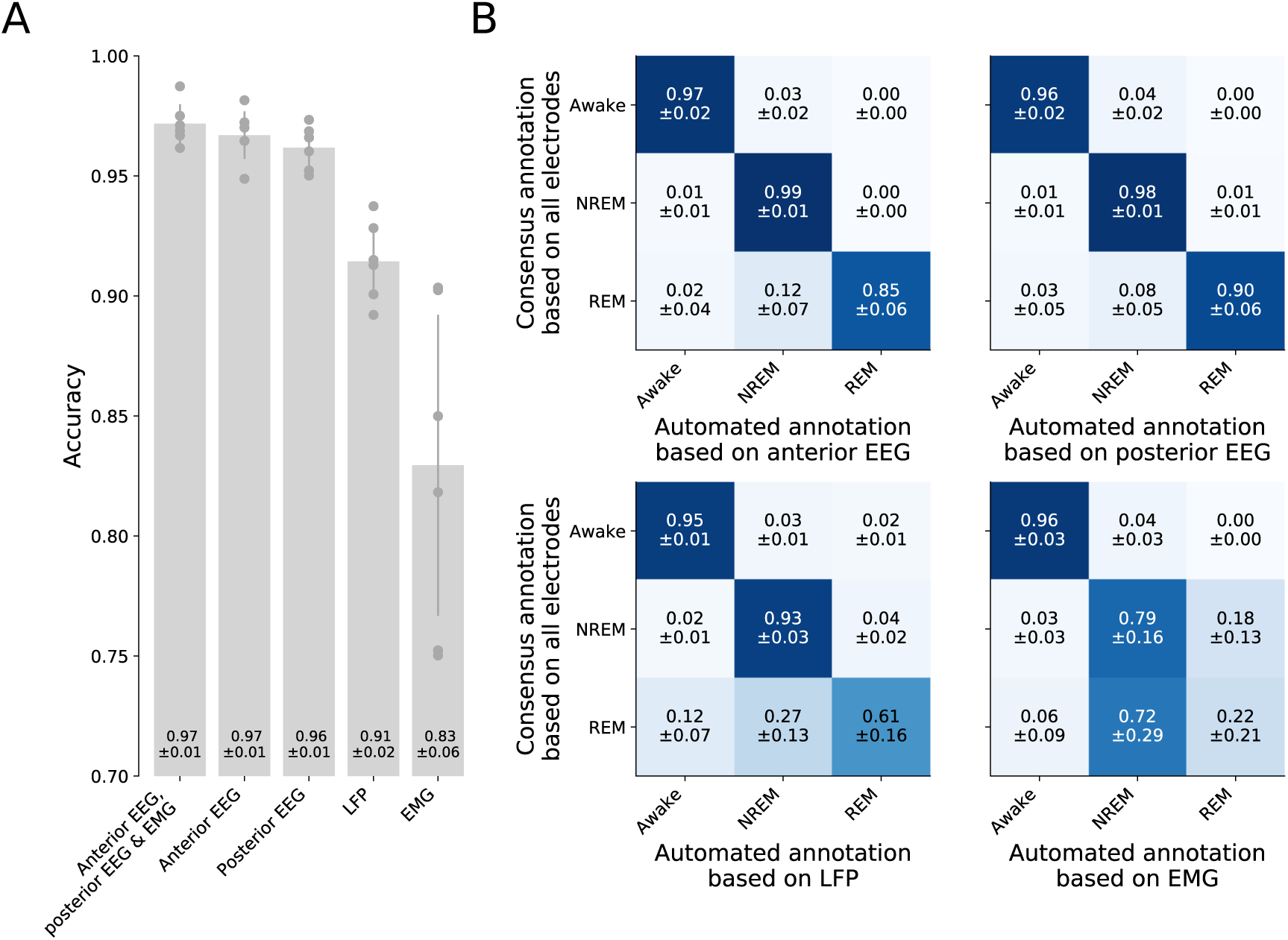
A single EEG signal is sufficient for Somnotate to infer vigilance states with high accuracy. (**A**) The accuracy of Somnotate’s sleep stage classification using a single input signal. Somnotate was trained and tested, in a hold-one-out fashion, on six 24-hour data sets. Only one signal was provided as an input signal: either the anterior EEG, the posterior EEG, the LFP from primary somatosensory cortex, or the EMG. (**B**) Confusion matrices when using only the anterior EEG (top left), the posterior EEG (top right), an LFP (bottom left), or the EMG (bottom right). Values indicate mean ± standard deviation. The overall accuracy of the predictions based on the anterior EEG alone did not differ from the overall accuracy when the anterior EEG, posterior EEG and EMG were provided simultaneously (p > 0.24, Wilcoxon signed rank test), as the small increase in the false negative detection rate for REM sleep was offset by an improved distinction between the awake and two sleep states. The overall accuracy of the predictions based on the posterior EEG was 1 % lower on average (p < 0.05, Wilcoxon signed rank test), although in this case the identification of REM showed a similar accuracy to when all signals were provided. The accuracy of predictions based on either an LFP recorded from primary somatosensory cortex, or only the EMG signal, was in both cases worse (by 6 % and 14 %, respectively), largely due to the performance on REM sleep episodes (p < 0.05 in both cases, Wilcoxon signed rank test). However, each individual signal was still sufficient to distinguish between awake and asleep states with high accuracy (∼95%), indicating that either signal would be sufficient in experiments that do not need to distinguish between REM and NREM sleep states.

**Supplementary Fig S5.**
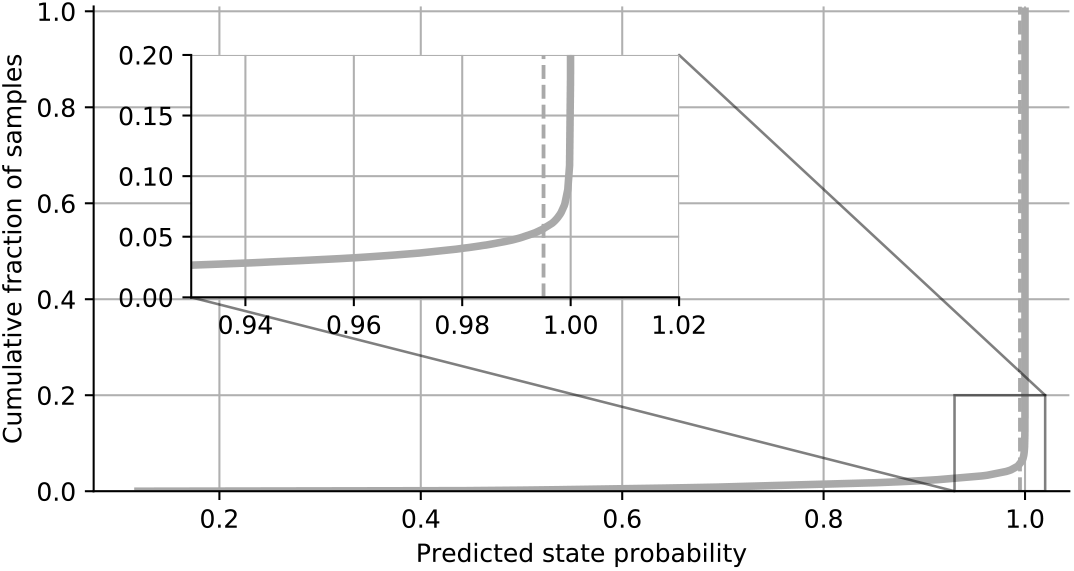
Selecting a state probability threshold to identify ambiguous samples. The probability of the predicted state was computed for each sample in six 24-hour data sets. As the cumulative distribution of probabilities exhibits an elbow at 0.995, this value was chosen as a threshold below which samples were classified as ambiguous.

**Supplementary Fig S6.**
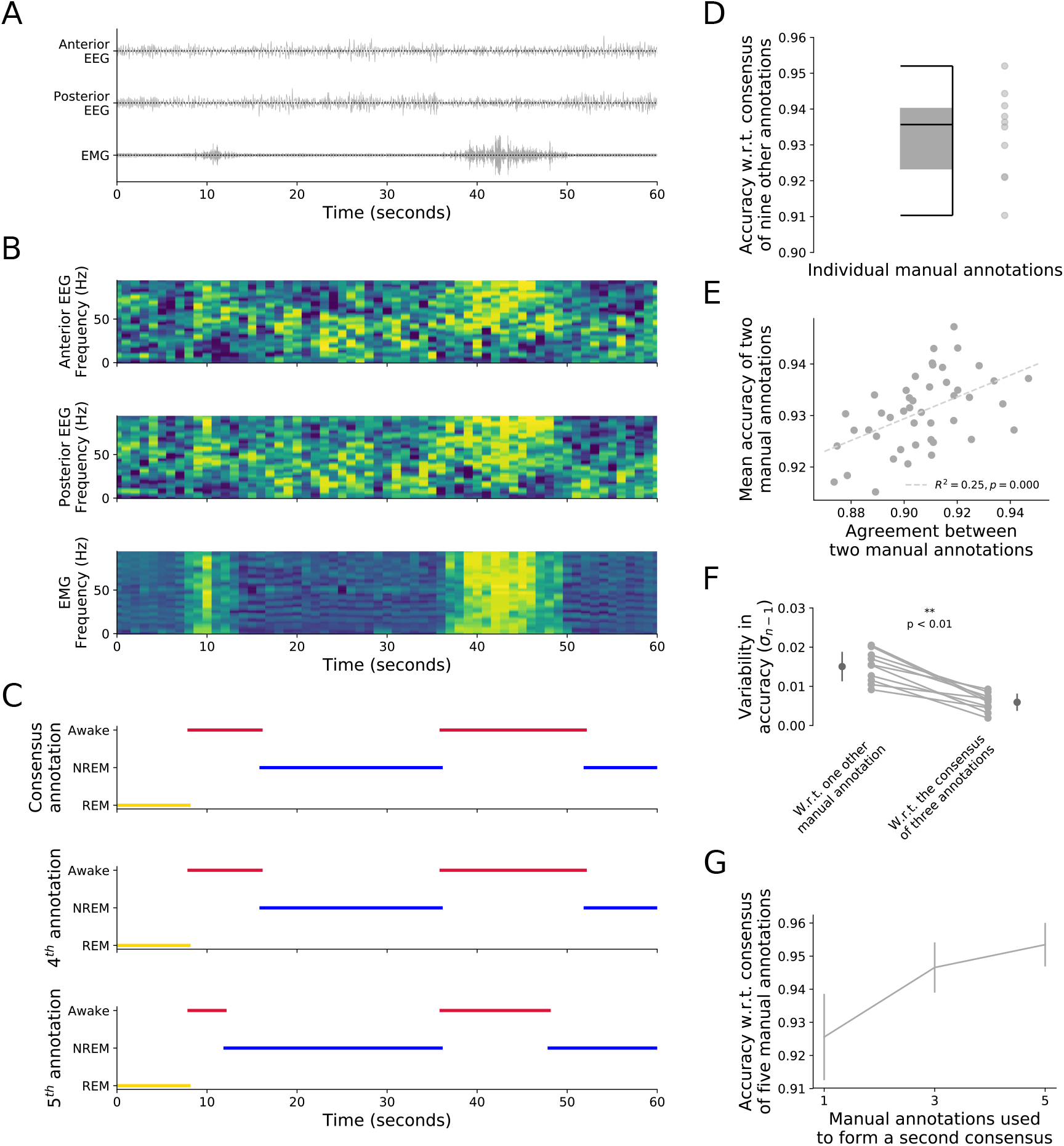
The consensus of manual annotations yields a better estimate of annotation accuracy. (**A**) The annotation of vigilance states was based on recordings of the anterior EEG, posterior EEG and EMG from a freely behaving mouse. A one-minute segment of the recordings is shown. (**B**) Multi-taper spectrograms for each of the recorded signals in ‘A’. (**C**) The majority-vote consensus of manual annotations by three independent experienced sleep researchers (top), which discriminates the vigilance states of ‘awake’ (red), ‘NREM’ sleep (blue) and ‘REM’ sleep (yellow). A fourth (middle) and fifth (bottom) independent individual manual annotation of the same segment. (**D**) A total of ten experienced sleep researchers independently annotated the same 12-hour recording and the accuracy of each annotation was assessed by using the consensus of the other nine annotations as a proxy for the ground truth. (**E**) For each possible pair of annotations, the inter-rater agreement was plotted against the mean accuracy of the pair of annotations, when judged against a consensus based on the remaining other eight annotations. (**F**) There was greater variability in the accuracy of an annotation when judged against a single other manual annotation, than when judged against the consensus of three randomly selected annotations (without replacement). Plot shows the variability in accuracy estimates (standard deviation with Bessel correction), which was significantly lower when using the consensus of three annotations (p < 0.01, Wilcoxon signed rank test). (**G**) A consensus was constructed from five of the ten independent annotations based on majority vote. A second consensus annotation was then constructed using either one, three or all five of the remaining annotations. The plot shows the mean agreement between the two consensus annotations. Error bars represent the standard deviation.

**Supplementary Fig S7.**
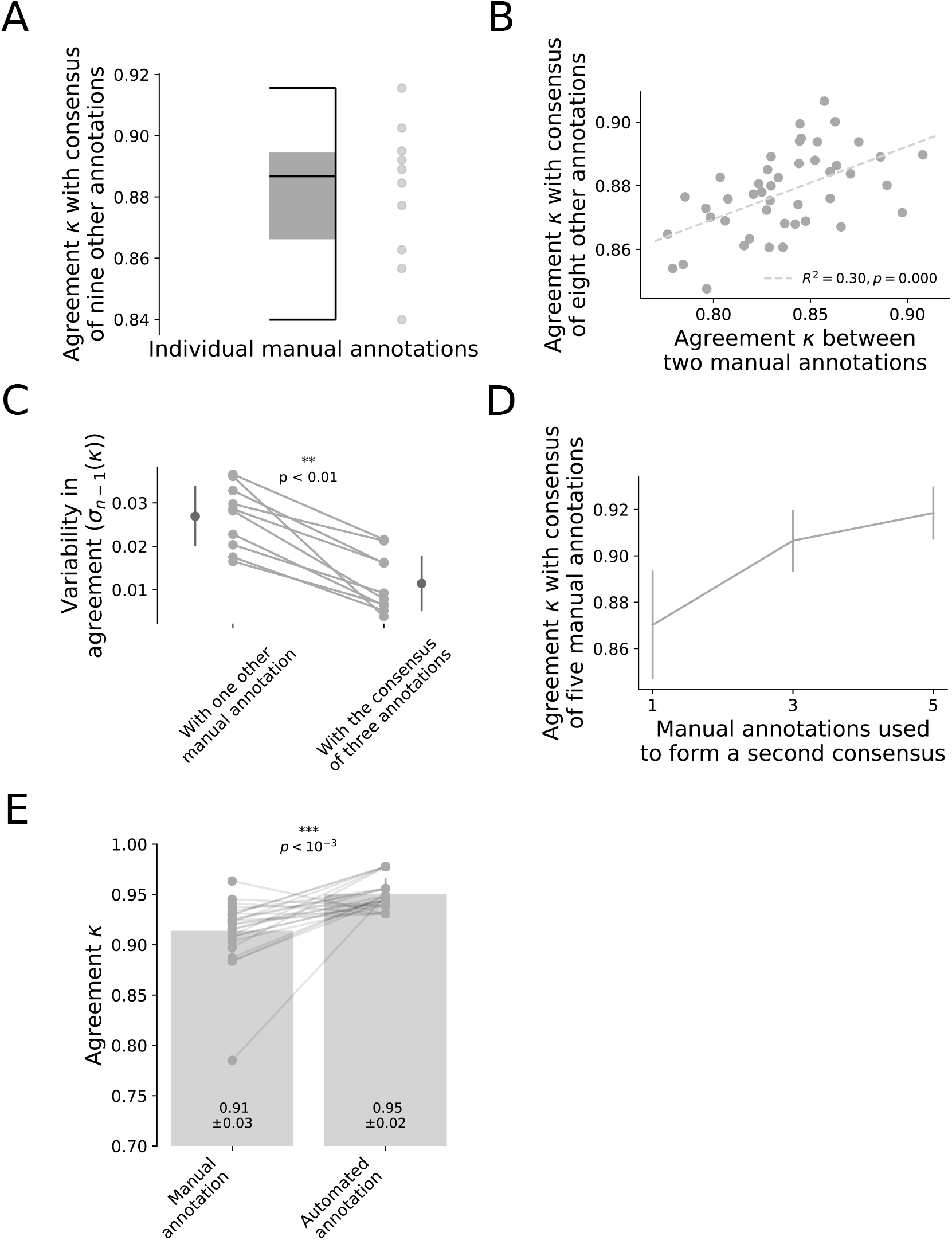
The consensus of manual annotations yields a better estimate of annotation accuracy, as measured by Cohen’s kappa. The analyses in Supplementary Fig S6D-G and Fig 2A were repeated using Cohen’s kappa as a measure of performance. (**A**) A total of ten experienced sleep researchers independently annotated the same 12-hour recording and the accuracy of each annotation was assessed by using the consensus of the other nine annotations as a proxy for the ground truth. (**B**) For each possible pair of annotations, the inter-rater agreement was plotted against the mean accuracy of the pair of annotations, when judged against a consensus based on the remaining other eight annotations. (**C**) There was greater variability in the accuracy of an annotation when judged against a single other manual annotation, than when judged against the consensus of three randomly selected annotations (without replacement). Plot shows the variability in accuracy estimates (standard deviation with Bessel correction), which was significantly lower when using the consensus of three annotations (p < 0.01, Wilcoxon signed rank test). (**D**) A consensus was constructed from five of the ten independent annotations based on majority vote. A second consensus annotation was then constructed using either one, three or all five of the remaining annotations. The plot shows the mean agreement between the two consensus annotations. Error bars represent standard deviation. (**E**) Somnotate was trained and tested, in a hold-one-out fashion, on six 24-hour data sets. Using a consensus annotation based on at least 3 manual annotations, the Cohen’s kappa score of the automated annotation was compared to the Cohen’s kappa score of individual manual annotations (n=25 manual annotations from 13 experienced sleep researchers). P-value is derived from a Wilcoxon signed rank test.

## Appendix S1

### Unbiased and precise assessment of automated and manual sleep annotation

The performance of sleep stage classifiers is typically measured by computing their agreement with two independent manual annotations. Performance is evaluated as the average agreement of the automated annotation with each of the two manual annotations, and this average is then compared to the level of agreement between the two manual annotations. This subtle difference in how manual and automated annotations are compared can lead to systematic biases in favour of the automated annotation. For example, assume that one manual annotation is perfectly accurate but the other manual annotation misclassifies half of the data. The inter-rater agreement between manual annotations is calculated as 1 * 0.5 = 0.5. Now assume that the automated annotation has exactly the average accuracy of the two manual annotations, i.e. 0.75. The average agreement with the two manual annotations will be (1 * 0.75 + 0.5 * 0.75) / 2 = 0.5625. In other words, the automated annotation will appear to be more than 10% better than the manual annotations, even though its accuracy was exactly average. Conversely, to achieve an average agreement score of 0.5, the automated annotation would only need to have an accuracy of 0.667, i.e. it could be 10% less accurate than the mean manual accuracy, while still achieving the same agreement between manual annotations.

For these reasons, we were keen to compare automated annotations to a majority-vote consensus derived from multiple independent manual annotations. We asked ten experienced sleep researchers (**Supplementary Table S1**) to annotate awake, NREM, and REM states from the same 12-hour data set based on simultaneous recordings of an anterior EEG, posterior EEG, and EMG in a freely behaving mouse (Materials and methods). This enabled us to generate consensus annotations based on multiple independent manual annotations (**Supplementary Fig S6A-C**). First, we assessed the accuracy of each annotation against the consensus of the other nine annotations. This revealed that although the overall accuracy of the annotations was high, individual annotations varied in terms of how closely they matched the consensus of the other annotations (**Supplementary Fig S6C-D**). This variance would cause systematic bias if one was to rely upon the level of agreement between just two manual annotations (see above). We were also keen to assess how precisely the agreement between any two annotators is able to capture the mean accuracy of both manual annotations. We therefore compared the inter-rater agreement for each pair of annotations to the mean of their accuracies based on the majority-vote consensus of the remaining eight annotations (serving as a proxy for ground truth). Whilst there was a statistically significant linear relationship between inter-rater agreement and the mean accuracy of the two annotations, the relationship was weak (R^2^ = 0.25; **Supplementary Fig S6E**).

To assess the quality of manual and automated annotations in an unbiased and more precise way, we compared the annotations to the consensus of multiple independent manual annotations. This comparison is unbiased as both the manual and automated annotations are assessed in exactly the same way. We confirmed that it is also a more precise measure, as the spread of performance estimates of manual annotations was smaller when using the consensus of three independent manual annotations to assess the accuracy of a fourth annotation, than when using a single other annotation as a point of reference (p < 0.01, Wilcoxon signed rank test; **Supplementary Fig S6F**). Finally, to estimate the minimum number of manual annotations required to achieve a high quality consensus sequence, we determined the consensus sequence of five annotations by majority-vote. Using either one, three or all five of the remaining unused annotations, we constructed a second consensus sequence, and computed the agreement between the two and then repeated this process for all possible combinations. On average, any individual manual annotation matched a consensus of five sequences for 92.5% ± 1.3% of the data (mean ± standard deviation), whereas a consensus of three annotations already significantly increased the agreement by 2.2% ± 1.5% (agreement 94.7% ± 0.8%; p < 0.01, Mann-Whitney rank test; **Supplementary Fig S6G**). There was a significant but more modest improvement of 0.5% ± 1.1%, when the number of manual annotations was increased to five (agreement 95.3% ± 0.7%; p < 0.01, Mann-Whitney rank test). Another widely used measure of inter-rater agreement is Cohen’s kappa, which accounts for the possibility of agreements occurring due to chance. When we repeated the analyses using this performance measure, we obtained analogous results (**Supplementary Fig S7)**.

In summary, a consensus derived from multiple independent manual annotations provides a less biased and more precise framework for assessing the quality of manual and automated annotations under comparable conditions. Based on these observations, we generated a larger test data set of six 24-hour EEG and EMG recordings (i.e. 144 hours total), which were independently scored by at least four experienced sleep researchers. This allowed us to compute the accuracy of manual and automated annotations using the majority-vote consensus of at least three other manual annotations for that recording. The recordings, individual manual annotations, and automated annotations are made freely available in standard formats at [WEBLINK TO PUBLIC DATA REPOSITORY].

## Notes

https://github.com/paulbrodersen/somnotate

### Competing Interest Statement

The authors have declared no competing interest.

### Summary of Updates

This version of the manuscript has been revised to conform to PLOS Comp Bio submission guidelines.

